# Clinical and Taxonomic Characterization of a Novel Urinary Tract Pathogen *Providencia lanzhouensis*

**DOI:** 10.1101/2025.01.14.632977

**Authors:** Yanghui Xiang, Xu Dong, Lan Ma, Dan Cao, Yi Li, Xiuzhi Jiang, Pusheng Xu, Xin Yuan, Kefan Bi, Yiru Zhang, Yuxin Han, Ying Zhang

## Abstract

**Background:** The genus *Providencia* includes species of ecological and clinical significance, with some acting as opportunistic pathogens in hospital-acquired infections such as urinary tract infection (UTI). However, overlapping phenotypic traits and genetic similarities pose challenges for accurate species characterization. This study exemplifies this challenge by identifying a novel *Providencia* isolate through genomic analysis.

**Results:** In this study, strain PAZ2 was isolated from the urine sample of a hospitalized patient in Lanzhou, Gansu Province, China. Initial mass spectrometry analysis identified the strain as *Providencia alcalifaciens*. It exhibits a resistance profile against antibiotics including ampicillin, tetracycline, tigecycline, polymyxin B, colistin, trimethoprim-sulfamethoxazole, and ciprofloxacin. However, whole-genome sequence analysis indicated that this strain is a novel species with an average nucleotide identity (ANI) values ranging from 79.92% to 94.73% and digital DNA-DNA hybridization (dDDH) values between 21.2% and 57.6% when compared with all known *Providencia* species. Furthermore, strain PAZ2 exhibited unique biochemical properties that differentiated it from all previously described species within the genus.

**Conclusions:** Genomic and biochemical evidence supports PAZ2 as a novel *Providencia* species, which we named *Providencia lanzhouensis* sp. nov. This finding enhances the taxonomy of *Providencia* and underscores its clinical relevance, notably in UTIs and other nosocomial infections. Accurate identification is crucial for guiding effective treatment and preventing misdiagnosis. Further studies should investigate its pathogenicity, antimicrobial resistance, and epidemiology to inform infection control measures.

## 1. Introduction

The *Providencia* genus comprises facultatively anaerobic, Gram-negative bacilli, originally classified within the *Enterobacteriaceae* family but subsequently reassigned to the *Morganellaceae* family following reclassification within the order *Enterobacterales* [1]. Species within the *Providencia* genus are widely distributed in diverse environmental settings, including water, soil, and various animal hosts [2]. These organisms are recognized as opportunistic pathogens, implicated in a range of clinical infections such as diarrhea, foodborne illnesses, urinary tract infections (UTIs), and bloodstream infections [3]. Currently, the genus *Providencia* includes 16 named species, *Providencia stuartii, Providencia manganoxydans, Providencia burhodogranariea, Providencia sneebia, Providencia rettgeri, Providencia huaxiensis, Providencia alcalifaciens, Providencia heimbachae, Providencia rustigianii, Providencia entomophila, Providencia wenzhouensis, Providencia xianensis, Providencia zhijiangensis and Providencia hangzhouensis* [4-13] and two novel recently identified species: *Providencia zhejiangensis* and *Providencia xihuensis* [14]. With *P. rettgeri, P. stuartii, P. alcalifaciens*, and *P. hangzhouensis* standing out due to their pronounced relevance to human infections. *P. stuartii* is notably associated with UTIs and healthcare-associated infections, particularly among patients with prolonged indwelling urinary catheters, thus representing a significant pathogen in nosocomial settings [15]. Another notable species, *P. rettgeri*, is also isolated from environmental sources like water and soil and has been linked to UTIs, diarrheal diseases, wound infections, and bacteremia [16]. This species presents a heightened infection risk for immunocompromised individuals and is frequently associated with secondary infections in burn patients.

In the 19th century, classification of *Providencia* species relied primarily on physiological characteristics and biochemical reactions. However, advances in molecular biology have led to the adoption of increasingly precise techniques for species identification and differentiation, particularly where biochemical profiles may overlap among species. Among these, 16S rRNA gene sequencing due to its high conservation across bacterial taxa has been instrumental in accurately identifying *Providencia* species, especially when conventional biochemical methods prove insufficient. Despite its utility, 16S rRNA sequencing has limitations in resolving intraspecies diversity [17]. Whole-genome sequencing (WGS), an advanced molecular tool, offers comprehensive insights into intraspecies variability and bacterial evolutionary patterns, establishing itself as the most robust method for accurate species identification.

In this study, we isolated a *Providencia* strain from a clinical sample collected at a tertiary hospital in Lanzhou. Initial identification using MALDI-TOF MS suggested the strain to be *P. alcalifaciens*. However, upon further analysis of distinct biochemical properties and whole-genome sequencing, we determined that this strain does not correspond to any known species within the *Providencia* genus. Therefore, we propose it as a novel species within the genus, tentatively named *Providencia lanzhouensis* sp. nov.

## 2. Materials and methods

### Strains and antibiotic susceptibility testing

Strain PAZ2 was isolated from the urine sample of a patient at a Lanzhou hospital as part of a routine clinical microbiology practice. Primary species identification of the strain was conducted using MALDI-TOF MS (bioMérieux). The antimicrobial susceptibility of the isolate was evaluated using agar dilution method to establish the minimum inhibitory concentrations (MICs), following protocols outlined in CLSI standards [18]. The interpretation of susceptibility breakpoints adhered to CLSI recommendations, with the exception of tigecycline assessment, which was conducted in accordance with EUCAST (European Committee on Antimicrobial Susceptibility Testing) parameters.

### Whole-genome sequencing and analysis

Bacterial DNA extraction was performed on this strain utilizing commercial DNA isolation reagents (QIAamp DNA Mini Kit manufactured by Qiagen). The genome sequencing strategy employed multiple platforms to ensure comprehensive coverage: short-read sequencing was conducted on an Illumina HiSeq 2500 platform to generate 150-bp paired-end reads, while long-read data was obtained using PacBio Sequel II technology. The integration of these complementary datasets was accomplished through the Unicycler pipeline [19] to construct complete genome assemblies. Taxonomic characterization involved comparative genomic analyses. Genome similarity metrics were calculated through two approaches: Average Nucleotide Identity (ANI) was computed using fastANI v1.32 [20], while digital DNA-DNA hybridization (dDDH) values were determined via the genome-to-genome distance calculator, implementing formula 2 [21]. Species boundaries were established based on accepted thresholds (ANI ≥96% and dDDH ≥70.0%) [22].

The genomic analysis pipeline included screening for antimicrobial resistance determinants using multiple specialized tools and databases: ABRicate (https://github.com/tseemann/abricate), AMRFinderPlus [23]. Genetic element identification and functional annotation were performed using the Bakta v1.9. 4 [24].

### Phylogenetic analysis

The ribosomal RNA analysis began with the amplification of 16S rRNA gene fragment by PCR sequencing using standard primers (27F/1492R) [25]. Reference sequences for comparative analysis were retrieved from EzBioCloud [26] and type strain genome repositories. Sequence organization and refinement involved MAFFT v7.505 [27] for initial alignment, followed by optimization with trimal v1.4 [28]. To enhance phylogenetic resolution, a multi-gene approach was implemented by analyzing five conserved housekeeping genes: *fusA* (encoding translational elongation factor EF-G), *gyrB* (DNA gyrase subunit B), *ileS* (isoleucyl-tRNA synthetase), *lepA* (translation elongation factor EF-4), and *leuS* (leucyl-tRNA synthetase). These sequences were extracted and combined from the study strains and relevant type strains, then processed through MAFFT and trimal pipelines. Genomic variation analysis was conducted through single nucleotide polymorphism (SNP) detection using the Snippy v4.16 (https://github.com/tseemann/snippy). Evolutionary relationships were reconstructed using maximum likelihood methodology in FastTree [29], with phylogenetic tree visualization accomplished through ggtree [30].

### Phenotypic characterization

As described in our previous study [16], Gram staining and biochemical characterization were performed using the bioMérieux API 20E and API 50CH systems, according to the manufacturer’s standardized protocols. Oxidase activity was assessed using bioMérieux’s oxidase test reagent. Cellular morphology was examined under an optical microscope following 16 hours of incubation on LB agar at 37°C. Growth characteristics were evaluated on various media, including tryptic soy agar, Luria-Bertani agar, blood heart infusion agar, and Müller-Hinton agar. Optimal growth conditions were systematically assessed in tryptic soy broth, varying temperature (4–44°C), pH (4.0–11.0), and salinity (0–9% w/v) within a controlled environment. Anaerobic growth potential was examined on BHI agar under anaerobic conditions for 48 hours. Catalase activity was determined by adding 3% hydrogen peroxide (v/v) to fresh bacterial biomass from LB agar cultures followed by observing for the presence of effervescence.

### Data availability

The complete genome sequence of PAZ2 from this study has been deposited in the GenBank database under BioProject accession PRJNA1180771.

## 3. Results

### 3.1 Clinical information and in vitro drug susceptibility test

The PAZ2 strain was isolated on 21 March 2024 from a urine sample of a patient. The patient presented with a history of dysuria with difficulty in urination persisting for over one month prior to hospital admission, attributed to prostatic hyperplasia, and was accompanied by urinary tract infection symptoms, including increased urination frequency and urgency. Strain PAZ2 was identified as *P. alcalifaciens* by MALDI-TOF MS, scoring 7.317 points. Following isolation and purification, the strain underwent comprehensive *in vitro* drug susceptibility testing. PAZ2 exhibited a narrow resistance profile, displaying resistance only to the antibiotics ampicillin, tetracycline, tigecycline, polymyxin B, colistin, trimethoprim-sulfamethoxazole and ciprofloxacin. Furthermore, the strain demonstrated susceptibility to a range of additional antibiotics, including ceftazidime, cefepime, imipenem, meropenem, amikacin, and gentamicin (see **Table 1**).

**Table 1.** Antibiotic susceptibilities of the strain PAZ2.

| Antibiotics | PAZ2 |  |
| --- | --- | --- |
|  | MIC (mg/L) | Category<sup>a</sup> |
| Ampicillin | >128 | R |
| Ceftazidime | <0.06 | S |
| Cefepime | <0.06 | S |
| Imipenem | 0.05 | S |
| Meropenem | <0.06 | S |
| Amikacin | 8 | S |
| Gentamicin | 2 | S |
| Aztreonam | <0.06 | S |
| Tetracycline | 64 | R |
| Tigecycline | 8 | R |
| Polymyxin B | 4 | R |
| Colistin | >128 | R |
| Trimethoprim-sulfamethoxazole | >128 | R |
| Ciprofloxacin | 32 | R |

<sup>a</sup>: S, susceptible; R, resistant.

### 3.2 Basic genomic features of strain PAZ2

To characterize this strain and its representative species, we performed whole-genome sequencing analysis of the strain PAZ2. The complete genome consists of a 4.12-Mb chromosome and a 2,683 bp plasmid, with an overall G + C content of 42.8% (**Fig. 1**). Genome analysis revealed 12 antimicrobial resistance genes, with 11 chromosomal and one plasmid-borne gene. The chromosomal resistance genes encode resistance to multiple antibiotic classes: aminoglycosides (*sat2, aadA1, aph(6)-Id, aph(3’’)-Ib, aph(3’)-Ia, aadA2*), sulfonamides (*sul2*), macrolides (*ere(A)*), trimethoprim (*dfrA32*), phenicols (*floR*), and tetracyclines (*tetC*). The plasmid-encoded *qnrD1* gene confers resistance to quinolones.

**Figure 1.**
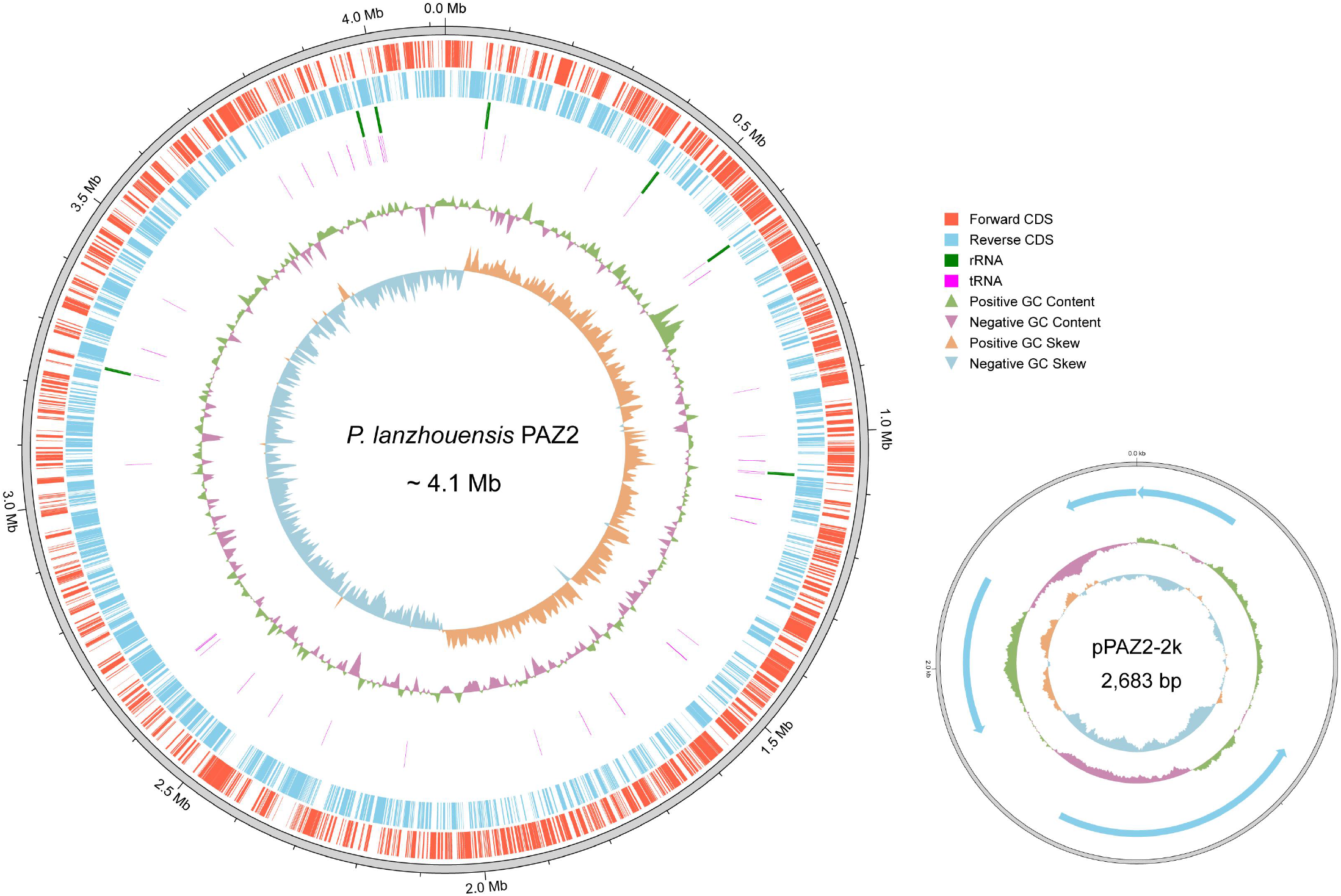
Structural diagrams of the chromosome and plasmid of PAZ2 identified in this study.

### 3.3 Taxonomic analysis reveals PAZ2 as a novel species distinct from known species

To establish the taxonomic position of strain PAZ2, we conducted a comprehensive phylogenetic analysis using multiple molecular approaches. Initial 16S rRNA gene sequence analysis revealed highest similarity (99.35%) with *P. huaxiensis* strain WCHPr000369 (NR_174258.1). Further phylogenetic analysis of 16S rRNA sequences from all known *Providencia* species indicated that PAZ2 showed closer evolutionary relationship to *P. stuartii* (**Fig. 2A**). However, given the known limitations of 16S rRNA-based classification for species delineation, we performed additional genomic analyses using ANI and dDDH.

**Figure 2.**
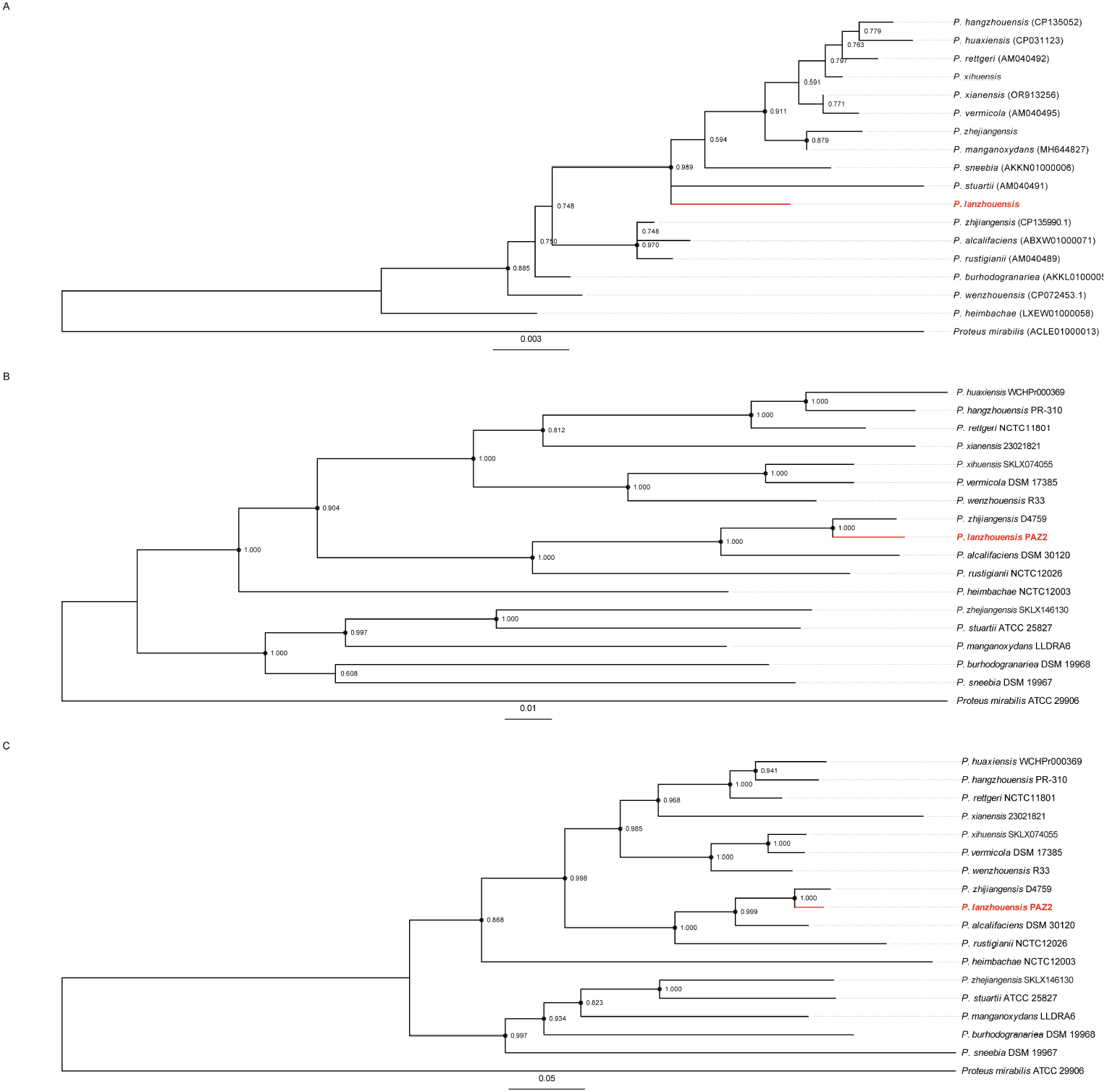
The evolutionary relationships within the *Providencia* genus were reconstructed through three distinct phylogenetic approaches. Panel (A) demonstrates the evolutionary framework inferred from 16S rRNA gene sequence analysis. Panel (B) represents phylogenetic associations derived from the concatenated sequences of five conserved housekeeping genes. Panel (C) illustrates the evolutionary patterns revealed through whole-genome SNP analysis. *Proteus mirabilis* ATCC 29906 serves as the designated outgroup species. The reliability of branching patterns was evaluated through 1,000 bootstrap replicates, with nodes displaying support values exceeding 50% being indicated. Notably, nodes with robust bootstrap support (>80%) are distinctly marked by solid black circles. Strains isolated and characterized in the present investigation are emphasized in red coloration.

Pairwise genome comparisons between PAZ2 and 16 type strains of known *Providencia* species yielded ANI values ranging from 79.92% to 94.73% and dDDH values from 21.2% to 57.6%, all below the established species threshold (ANI≥96%, dDDH≥70%). The highest ANI value (94.73%) was observed with *P. zhijiangensis*, suggesting a close but distinct relationship. Notably, comparison with previously unnamed taxonomic groups revealed that PAZ2 shared 97.10% ANI and 73.6% dDDH with taxon5, exceeding the species threshold and indicating that they belong to the same species (see **Table 2**).

**Table 2.** ANI and dDDH values comparing strain PAZ2 with know *Providencia* species.

| Species and/or strain | Assembly accession | PAZ2 |  |
| --- | --- | --- | --- |
|  |  | ANI (%) | dDDH (%) |
| *Providencia stuartii* ATCC 25827 | GCA_000154865.1 | 80.29 | 21.3 |
| *Providencia manganoxydans* LLDRA6 | GCA_016618195.1 | 80.52 | 21.9 |
| *Providencia burhodogranariae* DSM 19968 | GCA_000314855.2 | 79.92 | 21.1 |
| *Providencia sneebia* DSM 19967 | GCA_000314895.2 | 79.94 | 21.3 |
| *Providencia hangzhouensis* PR-310 | GCA_029193595.2 | 81.25 | 21.7 |
| *Providencia rettgeri* NCTC11801 | GCA_900455085.1 | 81.63 | 21.9 |
| *Providencia huaxiensis* WCHPr000369 | GCA_002843235.3 | 81.63 | 22.2 |
| *Providencia wenzhouensis* R33 | GCA_019343475.1 | 81.22 | 21.9 |
| *Providencia alcalifaciens* DSM 30120 | GCA_000173415.1 | 87.91 | 31.3 |
| *Providencia heimbachae* ATCC 35613 | GCA_001655055.1 | 80.84 | 21.3 |
| *Providencia vermicola* DSM 17385 | GCA_020381325.1 | 81.11 | 21.2 |
| *Providencia rustigianii* DSM 4541 | GCA_000156395.1 | 83.99 | 24.4 |
| *Providencia zhijiangensis* D4759 | GCA_030315915.2 | 94.73 | 57.6 |
| *Providencia xianensis* 23021821 | GCA_034661195.1 | 80.98 | 21.4 |
| *Providencia xihuensis* SKLX074055 | GCA_041953405.1 | 81.24 | 21.7 |
| *Providencia zhejiangensis* SKLX146130 | GCA_041953975.1 | 80.37 | 21.2 |

To further validate these findings, we constructed phylogenetic trees based on five housekeeping genes and core single nucleotide polymorphisms (SNPs). Both analyses consistently placed PAZ2 within the genus *Providencia*, clustering with *P. zhijiangensis* (**Fig 2B, C**). The substantial branch length separating PAZ2 from *P. zhijiangensis* supports their classification as distinct species, despite potentially sharing similar evolutionary trajectories.

### 3.4 Physiological and metabolic features

**Table 3** presents a comprehensive comparative analysis of the biochemical profiles of strain PAZ2 against other established *Providencia* species. To determine the optimal incubation conditions for strain PAZ2 growth, we ran a series of growth tests and bioanalyses. The strain PAZ2 was capable of growing on a variety of culture media, including TSA, LBA, BHIA, and MHA at a temperature of 37°C with ambient air condition. Colonies appeared circular, raised, yellow, opaque, and with a smooth texture (**Fig. 3**). The temperature range of the strain was determined to be 22–42 °C, exhibiting optimal growth at 35 and 37°C. The bacterial cells were capable of surviving within a pH range of 5 to 9, with an optimal growth observed at pH 6.0–7.0. For salt tolerance, the bacteria were able to thrive within the conditions of 0-6% (w/v) NaCl. Morphologically, the organism comprises of Gram-negative, mobile, and facultatively anaerobic rods. Notably, this strain PAZ2 lacked oxidase activity. In comparison to *P. zhijiangensis*, this strain exhibited weaker tolerance to environmental stresses, especially in terms of alkali resistance and salt tolerance.

**Table 3.** Biochemical characteristics of strain PAZ2 and type strains of other *Providencia* species^a^.

| Characteristic | PAZ2 | 1 | 2 | 3 | 4 | 5 | 6 | 7 | 8 | 9 | 10 | 11 | 12 | 13 | 14 | 15 | 16 | 17 | 18 | 19 |
| --- | --- | --- | --- | --- | --- | --- | --- | --- | --- | --- | --- | --- | --- | --- | --- | --- | --- | --- | --- | --- |
| **API 20E tests:** |  |  |  |  |  |  |  |  |  |  |  |  |  |  |  |  |  |  |  |  |
| β-Galactosidase | - | - | - | - | - | - | - | - | - | - | + | - | - | - | - | - | - | ND | - | - |
| L-Arginine dihydrolase | - | - | - | - | - | - | - | - | - | - | + | - | - | - | - | - | - | ND | - | - |
| Lysine decarboxylase | - | - | - | - | - | - | - | - | - | - | - | - | - | - | - | - | - | - | - | - |
| L-Ornithine decarboxylase | - | - | - | - | - | - | - | - | - | - | + | - | - | - | - | - | - | ND | - | - |
| Citrate utilization | + | + | - | - | + | + | - | - | - | + | + | - | + | + | + | + | + | + | + | + |
| H<sub>2</sub>S production | - | - | - | - | - | - | - | - | - | - | - | - | - | - | - | - | - | - | - | - |
| Urea hydrolysis | - | + | - | - | - | + | + | - | + | - | - | - | - | + | + | + | + | - | + | - |
| Deaminase | + | + | + | + | + | + | + | + | + | + | + | + | + | + | + | + | + | ND | + | + |
| Indole production | + | + | + | + | - | + | + | + | + | + | - | + | - | - | - | + | + | - | + | + |
| Acetoin production | - | - | - | - | - | - | + | - | - | - | + | - | + | - | - | + | - | - | - | - |
| Gelatinase | - | - | - | - | - | - | - | - | - | - | + | - | - | - | - | - | - | ND | - | - |
| D-Glucose | + | + | + | + | + | + | + | + | + | + | + | + | + | + | + | + | + | + | + | + |
| D-Mannitol | - | + | - | + | - | + | - | - | + | - | + | + | + | + | + | + | + | + | + | + |
| Inositol | - | + | - | + | - | + | + | - | - | + | + | + | - | + | + | + | + | - | + | + |
| D-Sorbitol | - | - | - | - | - | - | - | - | + | - | + | - | - | - | - | - | - | ND | - | - |
| L-Rhamnose | - | - | - | - | - | - | + | - | - | - | + | - | - | - | - | + | - | ND | + | - |
| Sucrose | - | + | - | - | - | - | - | - | - | + | + | - | + | - | - | - | - | ND | - | - |
| Melibiose | - | - | - | - | - | - | - | - | - | - | + | - | - | - | - | - | - | ND | - | - |
| Amygdalin | - | - | - | - | - | + | + | - | + | - | - | - | - | + | + | + | + | ND | - | - |
| L-Arabinose | - | - | - | + | - | - | - | - | - | - | + | + | - | - | - | - | - | ND | - | - |
| **API 50CHE tests:** |  |  |  |  |  |  |  |  |  |  |  |  |  |  |  |  |  |  |  |  |
| Aesculin | - | - | + | + | + | + | + | + | + | - | + | - | - | + | + | ND | ND | ND | + | - |
| Arbutin | - | - | - | + | + | + | - | - | + | - | + | - | - | + | + | ND | ND | ND | + | - |
| Cellobiose | - | - | - | - | - | - | - | - | - | - | + | - | - | - | - | ND | ND | ND | - | - |
| glycerol | - | + | - | - | - | - | + | - | - | + | + | - | + | + | + | ND | ND | - | + | + |
| 2-ketogluconate | - | - | - | - | - | + | - | - | - | - | + | + | - | - | - | ND | ND | ND | - | - |
| D-Lyxose | - | + | - | - | - | - | - | - | - | + | - | - | - | - | - | ND | ND | ND | - | + |
| salicin | - | - | - | + | - | + | - | - | + | - | + | - | - | + | + | ND | ND | + | + | - |
| D-Xylose | - | - | - | - | - | - | - | - | + | - | + | - | - | - | - | ND | ND | ND | - | - |

<sup>a</sup> Strains: 1, *P. manganoxydans* LLDRA6; 2, *P. alcalifaciens* DSM 30120; 3, *P. burhodogranariea* DSM 19968; 4, *P. heimbachae* DSM 3591; 5, *P. huaxiensis* KCTC 62577; 6, *P. rettgeri* DSM4542; 7, *P. rustigianii* DSM 4541; 8, *P. sneebia* DSM 19967; 9, *P. stuartii* DSM 4539; 10, *P. thailandensis* KCTC 23281; 11, *P. vermicola* DSM 17385; 12, *P. zhijiangensis* D4759; 13, *P. xianensis* 2302182; 14, *P. huashanensis* CRE-3FA-0001; 15, *P. hangzhouensis* PR-310; 16, *P. entomophila* IO-23; 17, *P. wenzhouensis* R33; 18, *P. xihuensis* SKLX74055; 19, *P. zhejiangensis* SKLX146130. ND, not determine

**Figure 3.**
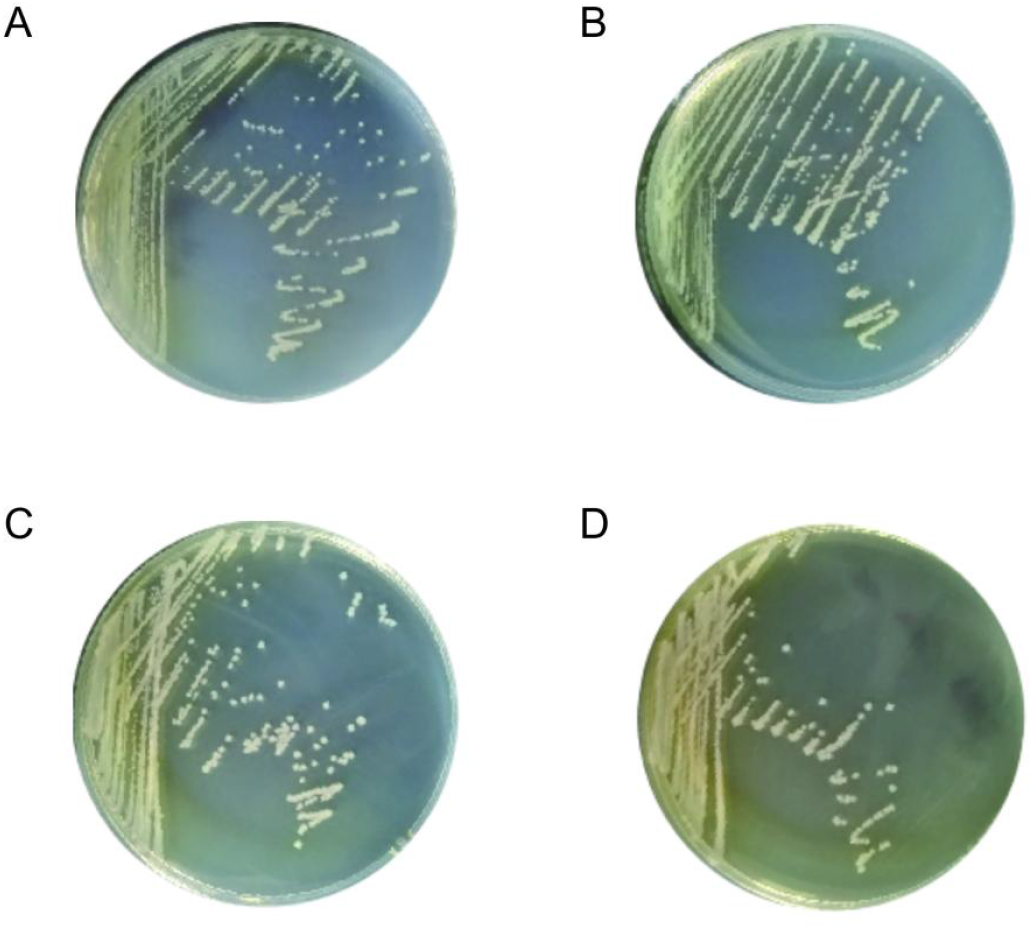
Morphological characteristics of PAZ2 The morphology of the strain PAZ2 in different culture media is shown in A-D. All bacteria were cultured at 37°C in an aerobic environment. A, Müller-Hinton Agar. B, Luria-Bertani agar. C, Trypticase soy agar. D, Brain Heart Infusion Agar.

### 3.5 Description of *Providencia. lanzhouensis* sp. nov

*P. lanzhouensis* (lan.zhou.en’sis. N.L. fem. adj. *lanzhouensis*, referring to Lanzhou City, Gansu Province, China, where the type strain was first isolated).

The cells are Gram-negative, motile, facultatively anaerobic rods, catalase-positive and oxidase-negative. The strain demonstrates robust growth on various culture media, including TSA, LB, BHI, and MH agar, with optimal growth observed at 37°C under aerobic conditions. Colonies are circular, raised, yellow, opaque, and smooth in texture. Growth occurs across a pH range of 5 to 9, with an optimal range between pH 6 and 7, and in NaCl concentrations ranging from 0 to 6% (w/v). Biochemical tests reveal positive results for deaminase activity and the fermentation of D-glucose, as well as positive results for citrate utilization.

The type strain, PAZ2 ^T^ =CCTCC AB 2024284^T^, was isolated from a urine specimen from Gansu Province, China. The DNA G + C content of the strain is 42.8%.

## 4. Discussion

Through mass spectrometry, we initially classified a clinical strain as belonging to the *Providencia* genus, but it was misidentified as an *P. alcalifaciens* species. However, through whole-genome sequencing analysis, we discovered that the species shares the highest ANI (94.73%) and dDDH (57.6%) values with *P. zhijiangensis*, although its ANI is still below the 96% threshold typically used to define a species, and its dDDH is well below the 70% threshold. This indicates that the strain does not belong to any known *Providencia* species but rather represents a new species within the genus. We also characterized its phenotypic properties and have named it *P. lanzhouensis* for the first time. The discovery of this new species contributes new data to bacterial taxonomy and evolution, aiding the development of a more precise bacterial classification system, while also helping to improve clinical microbiology databases for better pathogen identification.

Notably, similar misidentification by MALDI-TOF MS has been observed in the identification of *P. zhijiangensis* and *P. hangzhouensis* [12, 13]. These systematic errors highlight the limitations of current mass spectrometry-based identification systems, likely resulting from the rapid bacterial evolution and delayed updates of classification databases. This phenomenon emphasizes the necessity of integrating whole-genome sequencing and biochemical analyses for comprehensive identification in clinical diagnostics, while also underscoring the urgent need for developing more accurate identification systems.

*P. lanzhouensis* was isolated from the urine of an elderly male patient suffering from urinary retention. Its antimicrobial susceptibility profile shows that it remains relatively sensitive to most antibiotics, such as cephalosporins, carbapenems, and aminoglycosides, but is resistant to polymyxin and tetracycline antibiotics. Interestingly, this resistance pattern to polymyxin and tetracycline antibiotics is consistent with that observed in other *Providencia* species, although the underlying mechanism remains to be elucidated. While *P. lanzhouensis* shows close evolutionary relationship with *P. zhijiangensis*, they exhibit distinct biochemical characteristics, most notably in their inability to utilize D-mannitol and glycerol, which serve as crucial biochemical markers for species differentiation.

Our study has several limitations. Due to the limited sample size of *P. lanzhouensis*, the sample source is relatively singular, making it difficult to reveal its global or cross-regional distribution. Future studies should expand the sample collection to include more regions and clinical environments in order to better understand the distribution and epidemiology of this new species. Additionally, the clinical manifestations and pathogenicity of this new strain are not fully understood, especially with a limited sample size, which hinders the ability to assess its impact on different patient populations. Multi-center, interdisciplinary collaborative studies are necessary to accumulate more clinical cases and patient data, and conduct long-term epidemiological tracking to clarify its clinical features, pathogenicity, and infection mechanisms.

In short, the discovery of *P. lanzhouensis* not only enriches the taxonomy of the *Providencia* genus but also highlights potential deficiencies in the current bacterial identification systems, particularly the limitations of mass spectrometry in identifying new species. The antibiotic resistance features of this strain present new challenges for clinical treatment, underscoring the urgent need for further investigation into its resistance mechanisms, pathogenicity and its prevalence in clinical setting.

## Conclusions

This study highlights the intricate taxonomy of the *Providencia* genus and emphasizes the critical importance of accurate species identification for clinical pathogen recognition and effective treatment, particularly in the management of urinary tract infections (UTIs) and other hospital-acquired infections. Strain PAZ2, isolated from a hospitalized patient in Lanzhou, China, was initially misidentified as *P. alcalifaciens*. Comprehensive genomic and biochemical analyses, however, revealed its distinctiveness, with ANI and DDH values falling below species delineation thresholds and unique biochemical traits distinguishing it from known species. These findings support the designation of strain PAZ2 as a novel species, *Providencia lanzhouensis* sp. nov. This discovery not only enhances the understanding of *Providencia* diversity but also underscores its clinical relevance, as accurate identification is crucial for the timely and appropriate treatment of infections. Further studies are warranted to explore the pathogenic mechanisms, antimicrobial resistance profiles, and epidemiological patterns of *P. lanzhouensis*. Such research will provide critical insights into its potential role in UTIs and other infections, informing both clinical management strategies and infection control measures.

## Availability of data and materials

The complete genome sequence of PAZ2 from this study has been deposited in the GenBank database under BioProject accession PRJNA1180771. All other genomic data used in the present study are available at the public database.

## Abbreviations

*UTI*: Urinary Tract Infection
*MALDI-TOF MS*: Matrix-Assisted Laser Desorption/Ionization Time-of-Flight Mass Spectrometry
*ANI*: Average Nucleotide Identity
*dDDH*: Digital DNA–DNA Hybridization
*WGS*: Whole-genome Sequencing
*SNP*: Single Nucleotide Polymorphism
*CLSI*: Clinical & Laboratory Standards Institute
*EUCAST*: European Committee on Antimicrobial Susceptibility Testing
*MIC*: Minimum Inhibitory Concentration

## Acknowledgments

The authors would like to acknowledge all study participants who contributed to this study.

## Author contributions

YHX and YZ designed the study. YHX, LM, YL and XY collected the isolates and clinical data. XD analyzed, interpreted the data. YHX, DC, XZJ and PSX carried out the experiments. XD, KFB, YRZ and YXH prepared figures and visualizations for the manuscript. YHX and XD wrote the manuscript. All authors reviewed, revised and approved the final manuscript.

## Ethics declarations

### Ethics approval and consent to participate

Bacterial strains in this study were collected in clinical routine diagnostic lab practice, and ethical approval and patient consent were not required as no identifiable information was presented and in accordance with Decree No. 11 of the National Health Commission of the People’ s Republic of China (Operational Guideline for the Ethic Review of Biomedical Research Involving Human Subject, approved on September 30, 2016, implemented from December 1, 2016).

## Consent for publication

Not applicable.

